# Robust detection of tandem repeat expansions from long DNA reads

**DOI:** 10.1101/356931

**Authors:** Satomi Mitsuhashi, Martin C Frith, Takeshi Mizuguchi, Satoko Miyatake, Tomoko Toyota, Hiroaki Adachi, Yoko Oma, Yoshihiro Kino, Hiroaki Mitsuhashi, Naomichi Matsumoto

**Author notes:** These authors are contributed equally (Satomi Mitsuhashi, Martin C Frith). Corresponding authors: Martin C Frith, PhD Artificial Intelligence Research Center National Institute of Advanced Industrial Science and Technology (AIST) 2-3-26 Aomi, Koto-ku, Tokyo, 135-0064, Japan Satomi Mitsuhashi, MD, PhD Department of Human Genetics Yokohama City University Graduate School of Medicine Fukuura 3-9, Kanazawa-ku, Yokohama, 236-0004, Japan.

## Abstract

Tandemly repeated sequences are highly mutable and variable features of genomes. Tandem repeat expansions are responsible for a growing list of human diseases, even though it is hard to determine tandem repeat sequences with current DNA sequencing technology. Recent long-read technologies are promising, because the DNA reads are often longer than the repetitive regions, but are hampered by high error rates. Here, we report robust detection of human repeat expansions from careful alignments of long (PacBio and nanopore) reads to a reference genome. Our method (tandem-genotypes) is robust to systematic sequencing errors, inexact repeats with fuzzy boundaries, and low sequencing coverage. By comparing to healthy controls, we can prioritize pathological expansions within the top 10 out of 700000 tandem repeats in the genome. This may help to elucidate the many genetic diseases whose causes remain unknown.

## Introduction

A tandem repeat is a region where multiple adjacent copies of sequence reside in the genomic DNA. These regions are highly variable among individuals due to replication error during cell division. They are a source of phenotypic variability in disease and health. More than 30 human diseases are caused by copy-number alterations in tandem repeats^1^.

The range of pathogenic copy number change relative to the reference varies from a few to several thousand, and the length of repeating unit varies from e.g. three (triplet-repeat disease) to several thousand (macro-satellite repeat). As might be expected from such diverse underlying genetic causes, disease mechanisms are also variable. Well-known examples of triplet-repeat expansion diseases in protein-coding regions are polyglutamine diseases (e.g. spinal and bulbar muscular atrophy, Huntington’s disease)^2,3^. Triplet-repeat expansion of CAG or CAA codons, which encode glutamine, leads to toxic protein aggregation and neuronal cell death. Another example of triplet-repeat disease is caused by CUG repeat expansion in the 3’UTR of the transcript from the *DMPK* gene, producing a toxic gain-of-function transcript which sequesters splicing factor proteins and causes aberrant splicing, resulting in multiple symptoms^4^. Not only gain-of-function mutations, but also loss-of-function repeat change in the promoter region due to transcriptional silencing has been reported (e.g. fragile X syndrome)^5^. In addition to short tandem repeat diseases, repeat copy-number aberration in human disease is also reported in a macro-satellite repeat (D4Z4). Shortening of the D4Z4 repeat causes aberrant expression of the flanking gene *DUX4*, which has a toxic effect in muscle cells^6^. The thresholds of pathogenic repeat expansion in coding regions are usually less than 100 copies and sometimes even a few copy differences can cause disease (e.g. oculopharyngeal muscular dystrophy)^7^. In contrast, some disease causing tandem-repeat expansions in introns or UTRs can be very long (e.g. DMPK)^4^. Moreover, some repeats are interrupted by different sequences (e.g. *DMPK*, *ATXN10*, *SAMD12*)^8–10^, making it difficult to analyze the precise repeat structure.

It has been roughly a decade since the introduction of high throughput short read sequencers to clinical genetics. There have been numerous successful identifications of small nucleotide changes, especially in coding regions, mainly thanks to targeted sequencing (e.g. whole exome sequencing). However, the diagnostic rate remains 30% (depending on the diagnostic platform used)^11^, leaving a large population of Mendelian diseases unsolved. There may be many reasons, but one of the simplest is that the remaining patients may have mutations in “non-coding regions”, or they may have mutations in coding regions which were overlooked due to the limitations of short read sequencing. One candidate is tandem repeat regions, which are difficult to analyze genome-wide by conventional techniques. Identification of disease-causing tandem repeat changes is usually realized by classical genetic technologies (i.e. linkage analysis, Southern blot, etc.) and targeted repeat region analysis in a large number of families.

The recent advancement of long read sequencing technologies may provide a good solution, because long enough reads can encompass whole expanded repeats, and can be analyzed using the flanking unique sequences. Long read sequencers such as PacBio or nanopore sequencers have been coming to clinical genetics very recently. As of 2018, these technologies are continually improving in terms of accuracy and data output. However, in the clinical laboratory, it is still difficult due to cost-efficiency and the computing burden for large data. It would be preferable, and realistic, if low coverage data (~10X) can be used to detect alteration of tandem repeats.

We are aware of two previous methods for determining tandem repeat copy number from long DNA reads: PacmonSTR and RepeatHMM^12–13^. These methods: align the reads to a reference genome, then get the reads that cover a tandem repeat region of the reference, and perform sophisticated probability-based comparisons of these reads to the sequence of the repeating unit. In this study, however, we find that these methods do not always succeed with current long-read data.

We have recently advocated a method (using the LAST software) for aligning DNA reads to a genome allowing for rearrangements and duplications^14^. This method has two key features. First: it determines the rates of insertion, deletion, and each kind of substitution in the data, and uses these rates to determine the most probable alignments^15^. Second: it finds the most-probable division of each read into (one or more) parts together with the most probable alignment of each part. This method found diverse types of rearrangement, the most common of which was tandem multiplication (e.g. heptuplication), often of tandem-repeat regions^14^.

Here, we detect tandem repeat copy number changes by aligning long DNA reads to a reference genome with `last`, and analyzing these alignments in a crude-but-effective way. We point out several practical difficulties with analyzing tandem repeat sequences, which motivate our crude analysis method. Our approach is capable of analyzing tandem-repeats genome-wide even with relatively low coverage sequencing data. We believe that this tool will be very useful for identifying disease-causing mutations in tandem repeat regions which have been overlooked by short read sequencing in human disease.

## Methods

### tandem-genotypes method

`tandem-genotypes` (https://github.com/mcfrith/tandem-genotypes) has two required inputs: annotations of tandem repeats in a reference genome, and alignments of DNA reads to the same genome.

The annotations supply a start and end coordinate for each repeat, and the length *u* of its repeating unit. The repeat length need not be an integer multiple of the unit length. We define two ad hoc distances: “far”*f* = max[100, *u*] and “near”*n* = max[60, *u*]. (Actually, we truncate *f* at the edge of the sequence: where we speak of *f*, we really mean min[f distance to the edge of the reference sequence].)

`last-split` finds a division of each DNA read into (one or more) parts, and an alignment of each part to the genome. It gives each alignment a “mismap probability”, which is high if that part of the read aligns almost equally well to other loci^14^. We regard one read’s alignments as ordered by their 5’ to 3’ positions in the read.

For each DNA read, `tandem-genotypes` performs these steps:

1. Discard alignments with mismap probability > 10^-6^.
2. Discard alignments of mostly-lowercase sequence. This removes alignments that consist almost entirely of simple sequence (such as atatatatata), which are less reliable. Simple sequence is detected and lowercased by `lastdb` and `lastal`, using `tantan`^16^. An alignment is discarded if it lacks any segment with score >= `lastal`’s score threshold, when “gentle masking” is applied^17^.
3. Join consecutive alignments that are colinear on the same strand of the same chromosome, and separated by <=10^6^ bp.
4. Find all alignments that overlap a given repeat.

If there is one such alignment:

- Require that it extends beyond both sides of the repeat by at least *f*, else give up (i.e. don’t use this DNA read for this repeat).
- Extract all “gaps” from the alignment. Here, one “gap” may have *d* unaligned reference bases and *i* unaligned query bases, flanked by aligned bases. For each gap:

∘ If it does not overlap the repeat and *i* <= *u*/2: ignore it.
∘ If it is wholly >= *f* away from the repeat: ignore it.
∘ If it is partly >= *f* away from the repeat: give up.
∘ If it is wholly > *n* away from the repeat: ignore it.
∘ Define *r* to be the number of unaligned reference bases in the gap that are in the repeat.
∘ Define the gap’s net deletion size: *D* = min(*d - i*, *r*).
∘ Assume this gap contributes an integer (or zero) copy-number change. Find this contribution by rounding *D* to the nearest multiple of *u* (breaking ties by rounding towards zero).

If there is more than one such alignment:

- Require that they are consecutive in the read, and on the same strand.
- Define the “left” alignment to be the first one if the strand is “+”, else the last one. Define the “right” alignment in the opposite way.
- Require that the left alignment extends leftwards of the repeat, and leftwards of the other alignments, by at least *f*. Require that the other alignments do not extend leftwards of the repeat by *f* or more.
- Likewise for the right alignment.
- Define the insertion size as: the number of query bases, minus the number of reference bases (which could be negative), between the end of the left alignment and the start of the right alignment.
- Find the nearest multiple of *u* to this insertion size (as above).

### Prioritization of copy number changes

The repeats are ranked by a priority score. Each repeat has multiple predictions of copy-number change, one per DNA read. If the average number of predictions (for repeats with at least 1 prediction) is >= 3, ignore the most extreme expansion and contraction per repeat. For each repeat, take the most extreme remaining change, and calculate:

> (length increase in bases) / (reference repeat length + 30).

(The + 30 prevents excessive scores for short reference repeats.) This score is multiplied by an ad hoc value per gene annotation, currently: 50 for coding, 20 for UTR, 15 for promoter, 15 for exon of non-protein-coding RNA, and 5 for intron. This is multiplied by 2 for coding annotations where the repeating unit codes polyglutamine or polyalanine in any reading frame (out of 6). The repeats are ranked by absolute value of this priority score.

### Multi-dataset prioritization

Suppose we find tandem repeat changes in several individuals with and without a disease. For each repeat, calculate *d* = the cubic mean 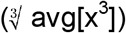 of the diseased individuals’ priority scores, and *h* = the cubic mean of the non-diseased individuals’ priority scores. If *d* < 0, negate *d* and *h*. Finally, the joint priority score is: max(*d* – max(*h*, 0), 0).

### Public and patient human genome data and alignment to reference genome

Human whole genome nanopore (rel3) and PacBio (SRR3197748) sequence data from the same individual (NA12878) were downloaded from (https://github.com/nanopore-wgs-consortium/NA12878) and from the SRA database, respectively^18^. Another 60X coverage human whole genome nanopore dataset from a different individual (NA19240) using PromethION was downloaded from https://www.ebi.ac.uk/ena/data/view/PRJEB26791.

Genomic DNA from a human patient with BAFME phenotype was sequenced by PacBio Sequel according to the manufacturer’s protocol. Briefly, genomic DNA was sheared by g-TUBE (Covaris, MA, USA), then size selection was done by BluePipin (Sage science, MA, USA) according to the standard method. These datasets were aligned to the human genome (GRCh38) with `last` version 936:

`windowmasker` -mk_counts -in genome.fa > genome.wmstat
`windowmasker` -ustat genome.wmstat -outfmt fasta -in genome.fa > genome-wm.fa
`lastdb` -P8 -uNEAR -R11 -c GRCh38 genome-wm.fa
`last-train` -P8 GRCh38 reads.fasta > train.out
`lastal` -P8 -p train.out GRCh38 reads.fasta | `last-split` > alns.maf□

### Generating tandem-repeat containing plasmids

Plasmids containing various numbers of CAG, GGGGCC and CAA used for this study were generated as described elsewhere^19–21^ and are available upon request (Supplemental Table1). Sequence data of the plasmids were deposited in (DRA007012, Table2).

A plasmid with CCTG repeats was generated as follows. Briefly, exon 1 with flanking 225 bp of intron 1 and exon 2 with flanking 1051 bp of intron 1 (including interrupted CCTG12 repeats) of the human *CNBP* gene were amplified by PCR using human genome DNA (cat.# G304A, Promega, Wisconsin, USA), and then cloned into pBluescriptII-KS(-) using In-Fusion cloning kit (Clontech Takara, Shiga, Japan). The resulting construct was digested with SalI and XhoI, and then ligated with T4 DNA ligase to delete the SalI site in the multicloning site of the pBluescriptII-KS(-). A new SalI enzyme site was generated by site-directed mutagenesis after 13bp of the interrupted CCTG_12_ repeats, to obtain a pBS-CNBP-SalI vector. Oligo DNAs containing CCTG15 repeats and flanking sequence, 5’-TCGA(CCTG)_15_C-3’ and 5’-TCGAC(CAGG)_15_-3’, were phosphorylated by T4 polynucleotide kinase, annealed, and then ligated using T4 DNA ligase (Takara, Shiga, Japan). The resulting ligated oligo DNA was digested with SalI and XhoI to remove undesired DNA fragments. Three tandemly ligated oligos were cloned into the SalI site of the pBS-CNBP-SalI vector. This plasmid has CCTG45 repeats interrupted by CTCGA in every 15 CCTG repeats, named interrupted CCTG_45_ (iCCTG_45_).

### Nanopore sequencing and alignment of tandem-repeat containing plasmids

These repeat-containing plasmids were digested with restriction enzymes NheI, EcoRI-HF, BamHI-HF or DraIII (NEB, MA, USA) (Supplemental Table 1) and then treated with Klenow Fragment DNA Polymerase (Takara, Shiga, Japan) at 37°C for 30 min. The whole DNA fragments were purified using AmPureXT beads (Agilent Techinologies, CA, USA), then subjected to nanopore sequencing. Library preparation was performed using a 1D Native barcoding genomic DNA kit (EXP-NBD103 and SQK-LSK108) then subjected to MinION (Oxford Nanopore Techonologies) sequencing using one FLA-MIN106(R9.4.1) flowcell according the manufacturer’s protocol. Base-calling and fastq conversion were performed with MinKNOW ver1.11.5. De-barcoding was done using EPIME software (Oxford Nanopore Techonologies).

Obtained fastq files were transformed to fasta files using seqkit fq2fa option (http://bioinf.shenwei.me/seqkit). fasta files were aligned to plasmid references like this;

`lastdb` -P8 -uNEAR -R01 plasmid-ref plasmid.fasta
`last-train` -P8 plasmid-ref reads.fasta > train.out
`lastal` -P8 -p train.out plasmid-ref reads.fasta | `last-split` > alns.maf□

### SCA10 data

SCA10 sequences from three patients with spinocerebellar ataxia 10 (MIM: 603516)^9^ were downloaded from SRA (subjectA: SRR2080459, subjectB: SRR2081063, subjectC-1: SRR2082412, subjectC-2: SRR2082428) then fasta files were generated by this command;

`fastq-dump` --fasta --table SEQUENCE --split-spot --skip-technical -I --gzip

Reads were aligned to the human genome (GRCh38) like this;

`lastdb` -P8 -uNEAR -R01 GRCh38□reference.fasta
`last-train` -P8 GRCh38 reads.fasta > train.out
`lastal` -P8 -p train.out GRCh38 reads.fasta | `last-split` > alns.maf□

### Comparison to RepeatHMM and PacmonSTR

The SCA10 and BAFME reads were analyzed as follows.

PacmonSTR:

`blasr` fasta reference.fa -m 5 --out blasr.out.m5
`makeBinnedAnchors.py` blasr.out.m5 simpleRepeat.txt 100
`pacMonStr_V1.py` blasr.out.m5 binned_anchors 10 8 current_directory_path

RepeatHMM:

`repeatHMM.py` FASTQinput --fastq fastq --Patternfile hg38.predefined.pa --repeatName gene --hgfile reference.fa
`repeatHMM.py` BAMinput --Onebamfile bam --Patternfile hg38.predefined.pa --repeatName gene --hgfile reference.fa

For SCA10 subjectA and subjectC-2, after consulting the RepeatHMM authors, we re-ran it with these options added: ‘--CompRep AlTlT50/C50lClT/C --SplitAndReAlign 1’.

### Running tandem-genotypes

The copy number changes of tandem repeats were determined by `tandem-genotypes` v.1.1.0 with some different options.

To check all tandem repeats in the human genome in NA12878 data (rel3 and SRR3197748); `tandem-genotypes` -u 1 -g refFlat.txt rmsk.txt alns.maf

To check disease-related tandem repeats in chimeric reads, BAFME and SCA10 data;

`tandem-genotypes` hg38-disease-tr.txt alns.maf

To check the plasmid sequences;

`tandem-genotypes` plasmid-repeat.bed alns.maf

Tandem repeat (rmsk.txt) and gene (refFlat.txt) annotations were obtained from the UCSC genome database (http://genome.ucsc.edu/)^22^. We made the file hg38-disease-tr.txt, with 31 disease-associated tandem repeats, based on Tang *et al*.^1^.

Tandem repeat changes in a chimpanzee, relative to the reference human genome, were found like this:

`tandem-genotypes` -g refFlat.txt rmsk.txt hg38-panTro5-1.maf These alignments (from https://github.com/mcfrith/last-genome-alignments)^14^ are of an assembled chimp genome, *not* long reads: our methods work in this case too.

Multi-dataset prioritization was done with commands of this form: `tandem-genotypes-join` data1 : data2

data1: `tandem-genotypes` output to be prioritized

data2: `tandem-genotypes` output to be de-prioritized

For the BAFME patient, two humans and one chimpanzee (NA12878 PacBio SRR3197748, NA19240 PromethION ERR2585112-5, and panTro5) were used for de-prioritizing possibly benign expansions. For rel3 data with coding-region expansions, NA19240 (PromethION ERR2585112-5) the BAFME patient (PacBio), and panTro5 were used for de-prioritization

### Chimeric reads of plasmid-derived repeats and human-derived flanks

Human nanopore reads covering repeat expansion disease loci were extracted from whole genome sequence data (rel3). The nanopore sequences flanking the repeat were excised using an in-house script. Randomly selected expanded and non-expanded repeat sequences were excised from plasmid nanopore sequences, using `maf-cut`. Then, these expanded and non-expanded repeats were inserted between the flanks, to generate chimeric reads. The combinations of repeat copy and number of rel3 reads are shown in Table 1, imitating the diploid genome. The chimeric reads were aligned to GRCh38 as mentioned above with `WindowMasker`^23^.

Dot-plot pictures were made with last-dotplot using the following command;

`last-dotplot` --max-gap2=0,inf --rmsk1 rmsk.txt aln.maf file.png

### PCR amplification of inexact repeats in rel3 and PacBio data

Two inexact tandem-repeats were tested by PCR and Sanger sequencing. Primers are described in Supplemental Table 2. PCR amplification was done using KAPA HiFi HS DNA polymerase (Kapa Biosystems, Basel, Switzerland). DNA for NA12878 was obtained from the Coriell Institute (https://coriell.org). PCR products were cloned into pCR-Blunt vector (Thermo Fisher Scientific, MA, USA) and subjected to Sanger sequencing.

### Ethics

The Institutional Review Board of Yokohama City University of Medicine approved the experimental protocols. Informed consent was obtained from the patient, in accordance with Japanese regulatory requirements.

## Results

### Nanopore sequencing of tandem-repeat containing plasmids

We tested plasmids with four different kinds of repeat (CAG, CAA, GGGGCC and iCCTG) that are known to cause human diseases. The CAG plasmids have 5 different copy numbers (6, 18, 30, 70 and 130). The CAA and GGGGCC plasmids have two different copy numbers, 15/109 and 21/52, respectively. These plasmids were subjected to Oxford Nanopore Technologies’ (ONT) MinION sequencing. The MinION reads were aligned to plasmid reference sequences with copy number 6 (CAG), 15 (CAA), 3 (GGGGCC), and 15 (CCTG). `tandem-genotypes` predicts the copy number change in each read: these predictions roughly agree with the actual copy number changes (Figure 1, red arrows). There is a minority of unexpectedly low copy number predictions (Fig1d,i; arrow), especially for the longer plasmids: manual inspection of alignment dotplots (not shown) suggests that these are correct, and the copy numbers in these plasmids may not be completely stable.

**Figure 1.**
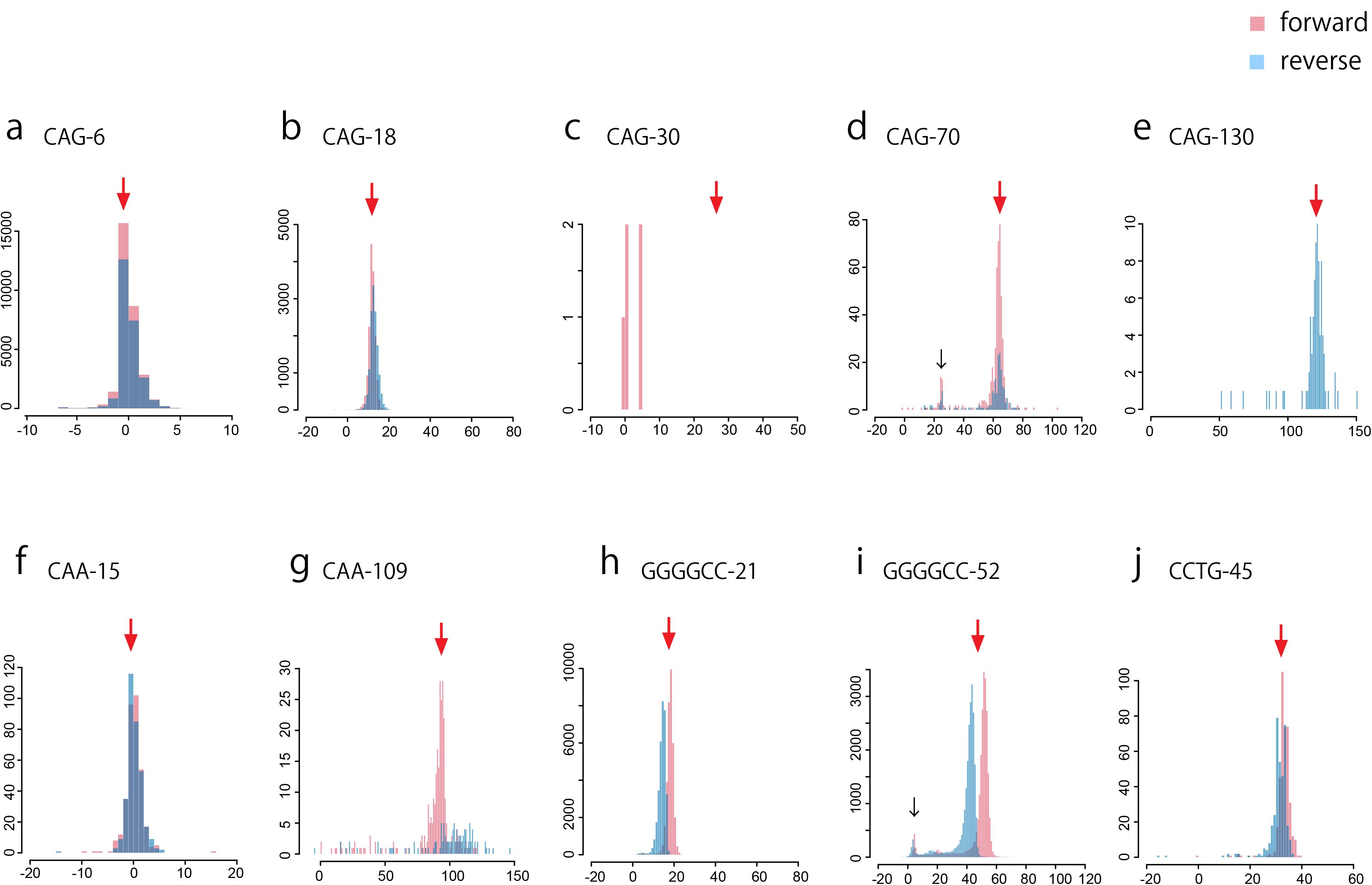
Distribution of predicted change in repeat copy number, for nanopore reads from each of ten plasmids. Red arrows: projected copy number change. Forward (red) and reverse strand reads (blue) are shown separately. y-axis: read count, x-axis: change in copy number relative to the reference plasmid. Reference copy numbers in each plasmid are in Supplemental Table 1. Black arrows: reads in these peaks may actually have shortened repeats.

In one case, pBS-(CAG)30, `tandem-genotypes` fails with almost no predictions. pBS-(CAG)30 was digested at an enzyme site very near the repeat region (10-bp upstream), so there is only 10 bp of non-repeat sequence upstream of the repeat, which is too short for step 2 of `tandem-genotypes`. Thus, we digested the same plasmid with a different enzyme, DraIII, far from the start of the repeat. As expected, the `tandem-genotypes` prediction agrees with the actual copy number change (Supplemental Figure 1a, red arrows).

The GGGGCC repeats have bimodal copy number predictions, where the two modes correspond to reads from each DNA strand (Fig 1h-i, Supplemental Fig 1b). This can be explained by sequence-specific sequencing error tendencies^24^ that differ for the forward- and reverse-strand repeat sequences.

In human genomic sequencing, it might be difficult to obtain deep coverage such as 1000X as we have done for these plasmids, due to cost. For some repeat expansion diseases, such as polyglutamine disease, disease-causing copy number change thresholds from the controls are usually less than 100. To test the ability of `tandem-genotypes` to distinguish fine copy number differences of these plasmids even with low coverage, we randomly picked 50, 30 and 15 reads from each dataset and compared the copy number predictions. Even with low coverage (15X) it is not difficult to distinguish copy numbers 18, 30, 70 and 130 for CAG repeats; 15 and 109 for CAA repeats; 21 and 52 for GGGGCC repeats (Figure 2).

**Figure 2.**
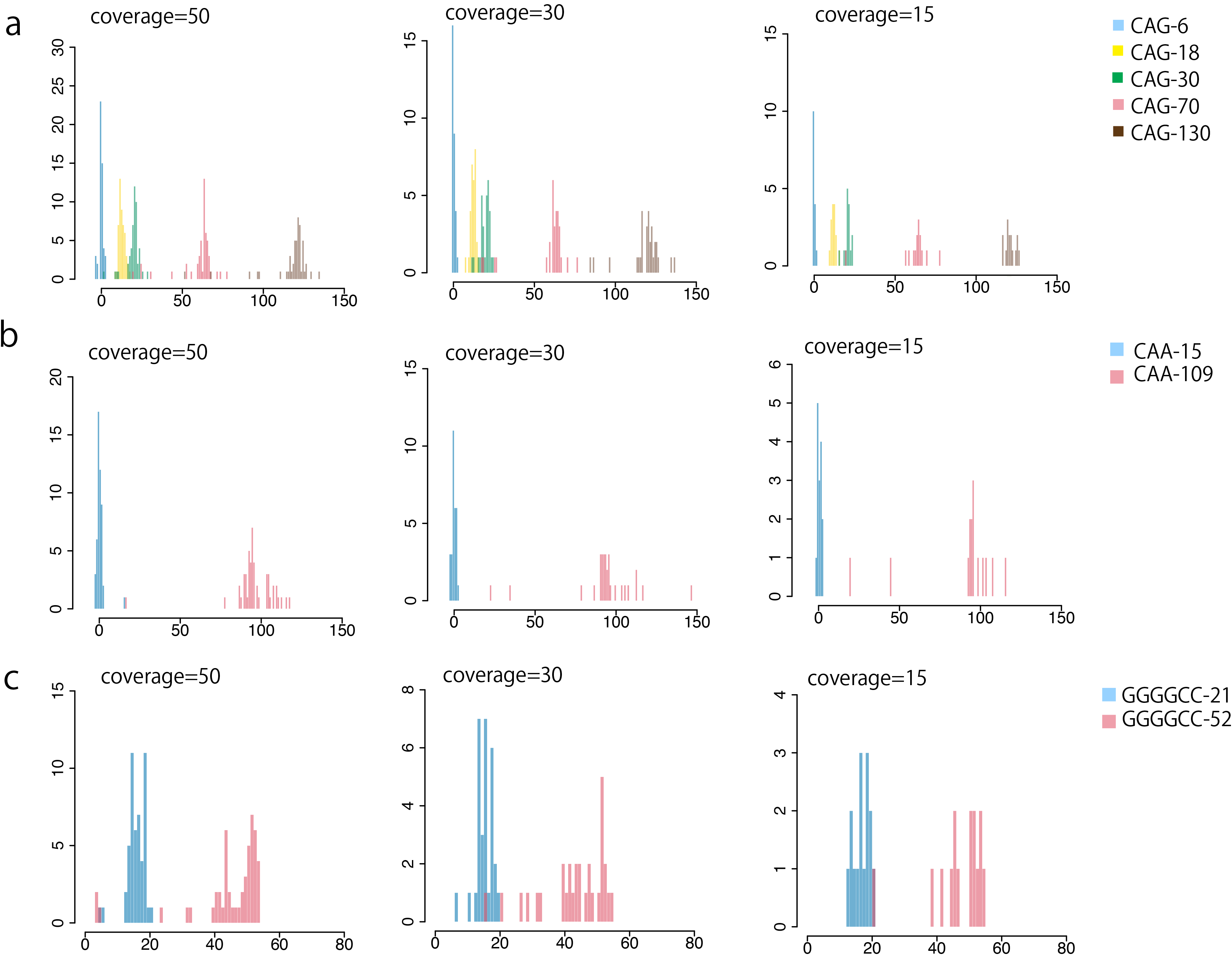
Distribution of predicted change in repeat copy number, for random samples of plasmid nanopore reads. Coverages 50, 30 and 15 were tested. (a) CAG repeats. (b) CAA repeats. (c) GGGGCC repeats. y-axis: read count, x-axis: change in copy number relative to the reference plasmid.

### Analyzing chimeric human/plasmid nanopore reads

We performed further tests on semi-artificial data. We obtained human nanopore reads that cover 10 disease-associated repeat regions, and replaced the repeat region in each read with the repeat region of a plasmid nanopore read. We used plasmid repeat regions with disease-causing and healthy repeat copy numbers in 1:1 ratio (Table1). These chimeric reads were aligned to a reference human genome, and copy-number changes were predicted. For each repeat, the predictions have clear bimodal distributions (Figure 3). As these plasmid-origin repeats were expected to vary because of nanopore base-call error or replication error in *Escherichia coli* during plasmid preparation, we also made chimeric sequences by inserting the exact number of exact repeats, to test the accuracy of `tandem-genotypes`. The results were improved in all cases (Figure 3), in particular the *C9orf72* case with large strand-bias, indicating that deviations from the projected copy numbers are due to nanopore sequencing errors or replication error, and not systematic errors of `tandem-genotypes`.

**Figure 3.**
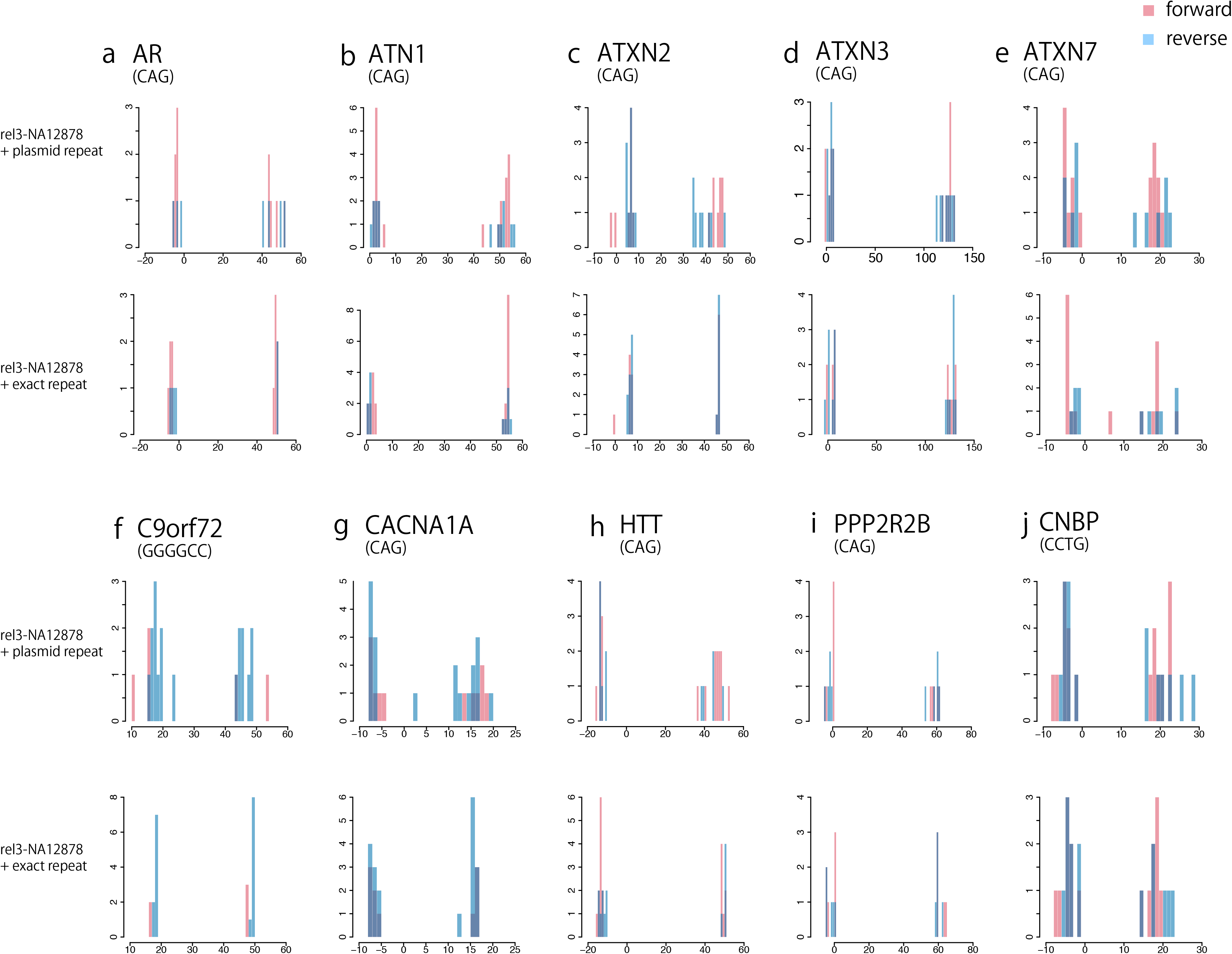
Distribution of predicted change in repeat copy number, for nanopore reads of human DNA with inserted repeats. Reads covering each of ten disease-associated repeat loci were selected, and the repeat region in each read was replaced by: the repeat region of a plasmid nanopore read (a-j top panels), or exact repeats (a-j bottom panels). y-axis: read count, x-axis: change in copy number relative to the reference human genome. Forward (red) and reverse strand reads (blue) are shown separately.

### Pacbio sequencing datasets of patients with SCA10

Next, we examined four PacBio sequencing datasets of cloned PCR amplification products from the SCA10 disease locus (spinocerebellar ataxia 10, MIM 603516). SCA10 is caused by ATTCT expansion in the intron of *ATXN10*.These datasets are from three unrelated patients: A, B and C^9^. Patient C has two datasets, C-1 and C-2, which are different clones sequenced with different PacBio chemistries. According to McFarland et al., subjects A, B and C can be estimated to have 4.5 kb, 3.9 kb and 2.7 kb repeat expansions (since PCR product sizes are 6.5 kb, 5.9 kb and 4.7 kb and they contain 2 kb flanking sequences), respectively^9^. `tandem-genotypes` predicted datasets A, B, C-1 and C-2 have average expansions of 913, 841, 484 and 486 copies relative to the reference (14 copies), hence repeat lengths 4.6 kb, 4.3 kb, 2.5 kb and 2.5 kb respectively (Figure 4a). Thus subject B is predicted to have 0.4 kb larger repeat size than the PCR product, however from McFarland et al. Figure 1b^9^, the purified clone they sequenced by PacBio had > 6 kb insertion, making the actual repeat size > 4 kb, closer to our prediction.

**Figure 4.**
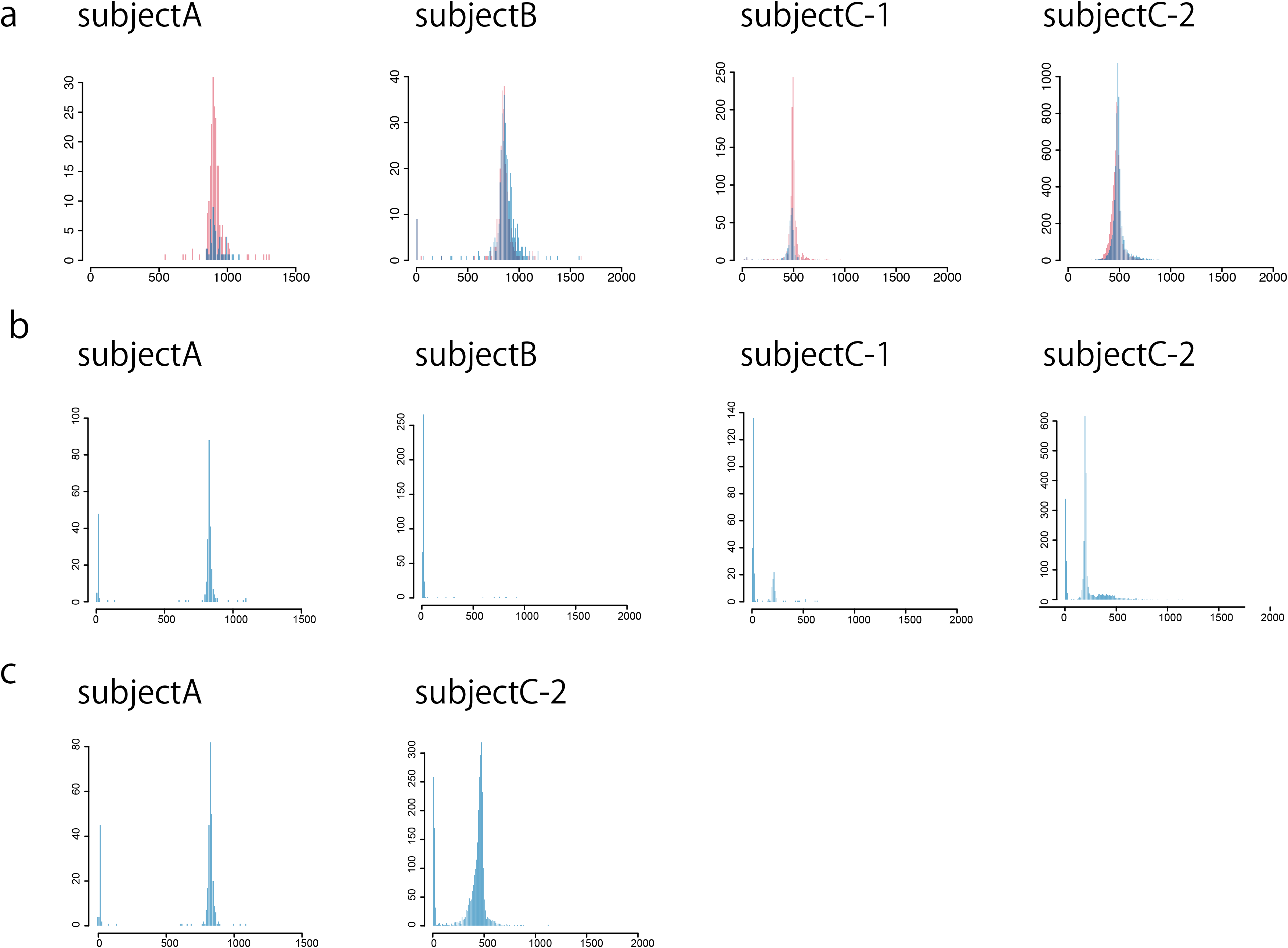
Distribution of predicted change in repeat copy number, for PacBio (RSII) reads of cloned SCA10 loci from 3 patients. (a) `tandem-genotypes`. Forward (red) and reverse strand reads (blue) are shown separately. (b) RepeatHMM with straightforward parameters. (c) RepeatHMM with non-obvious parameters suggested by its authors. y-axis: read count, x-axis: change in copy number relative to the reference human genome.

The same datasets were also analyzed with RepeatHMM and PacmonSTR. We first ran RepeatHMM with straightforward parameters: for subject A it found a similar peak to us but also an unexpected peak around zero, and it did not find the expected peaks for the other three datasets (Figure 4b). We then consulted the RepeatHMM authors, who suggested non-obvious parameters that improved the C-2 result, but there was still a peak around zero (Figure 4c). PacmonSTR did not detect the expected repeat numbers (data not shown).

### PCR and Sanger sequencing of non-exact expanded repeats in NA12878

Repeat annotations (i.e. rmsk from UCSC) include non-exact tandem-repeats. Non-exact or interrupted tandem-repeats sometimes cause human disease^8^. We detected inexact repeat expansions in the NA12878 human genome, by applying `tandem-genotypes` to the PacBio and MinION datasets.

In an intron of *PCDH15*, rmsk annotates a “TATAT” tandem repeat (chr10:54421448-54421530), though the actual sequence is not exactly TATAT (tatataaaataaactttattatatttagcatttgatttttatttatgtatattataaaatgaatatagtttatattataata). `tandem-genotypes` found two peaks for this repeat, indicating a heterozygous ~300 bp insertion (Figure 5a). PCR amplification of this region showed two different products possibly from different alleles. One had the same length as the reference sequence (PCDH15-intron-repeat-S) (Figure 5a). Sanger sequence of the other longer PCR product (PCDH15-intron-repeat-L) revealed a ~332 bp insertion. Surprisingly, this insertion was not a tandem expansion, but rather an AluYb8 SINE (according to RepeatMasker).

**Figure 5.**
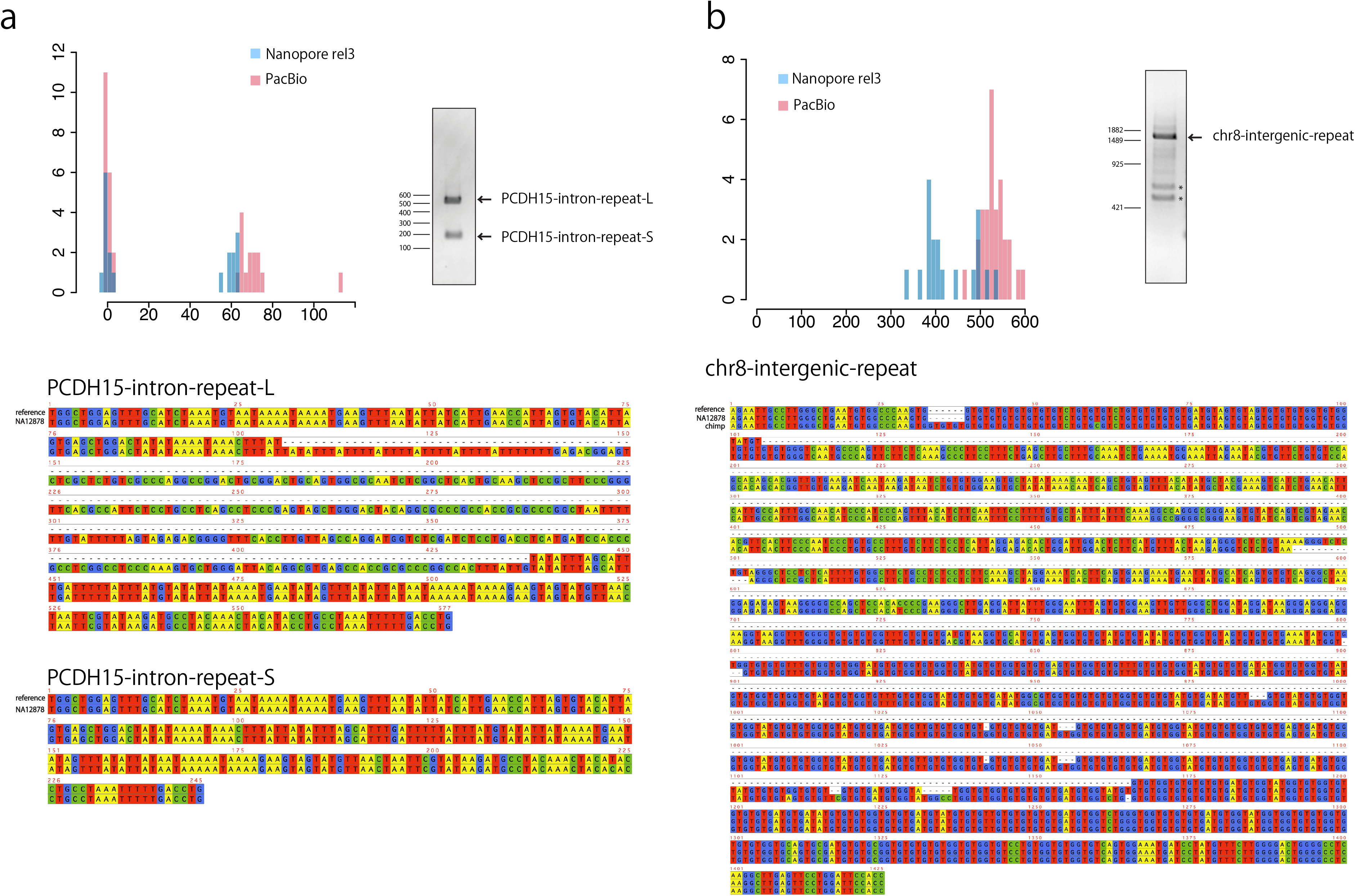
PCR results and Sanger sequencing of two tandem-repeat loci in human genome NA12878, with expansions relative to the reference human genome (hg38). The histograms show read counts (y-axis) for predicted copy number changes (x-axis), with nanopore (rel3) in blue and PacBio (SRR3197748) in red (a) Expansion of an inexact TATAT repeat in an intron of *PCDH15:* actually insertion of an AluYb8 SINE. (b) Expansion of an intergenic GT repeat: actually deletion in the reference human genome. The bands marked by * were sequenced and proved to be non-target amplification.

We also examined an intergenic GT tandem repeat (chr8:48173947-48174212). `tandem-genotypes` found one peak indicating an insertion of ~1000 bp (Figure 5b). PCR of this region showed a single product from this tandem repeat region, estimated to contain a ~1000 bp insertion. Sanger sequencing revealed that this expansion includes not only GT but also some unknown sequence. The expanded sequence is present in the chimpanzee genome (Figure 5b), indicating that this is actually a deletion in the human reference genome (which may have occurred by recombination between GT tandem repeats).

These two examples indicate that `tandem-genotypes` can also find complex and interrupted expansions (or non-deletions) of tandem repeats.

### Pacbio sequencing of a patient with BAFME

We further analyzed PacBio whole genome sequencing of a patient with a phenotype of benign adult familial myoclonic epilepsy (BAFME). In another large number of BAFME patients in Japan^10^, the cause was attributed to large expansions of intronic TTTCA and TTTTA repeats in *SAMD12*. We wished to know whether our patient has such an expansion in *SAMD12*. We sequenced this patient’s genomic DNA using a PacBio Sequel sequencer. `tandem-genotypes` detected ~5 kb insertion in three reads at the BAFME locus, where the coverage is 6X (Figure 6a). We also applied RepeatHMM and PacmonSTR to this dataset, but they failed to predict any expansions.

**Figure 6.**
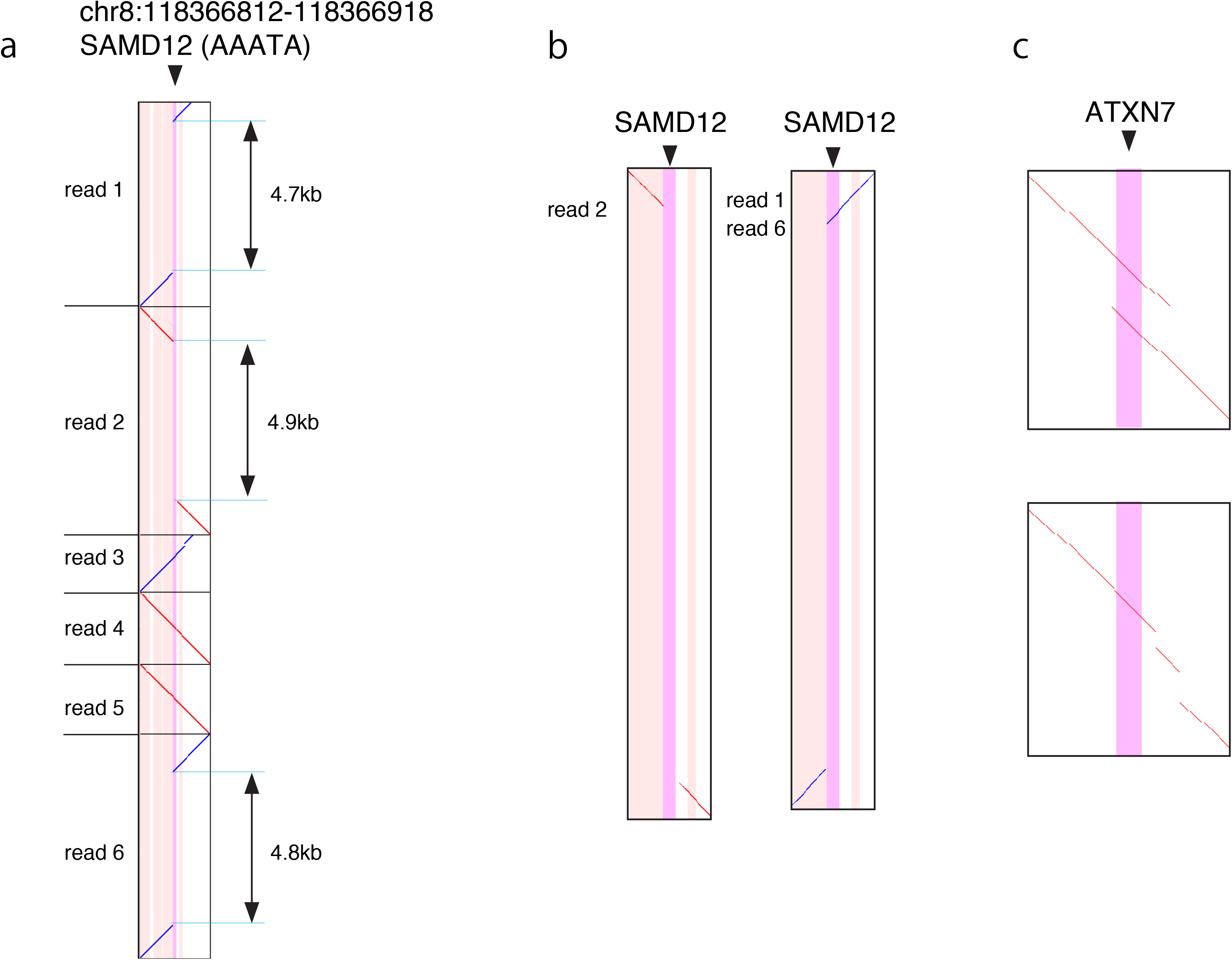
Alignments of DNA reads (vertical) to the reference human genome (horizontal). Diagonal lines indicate alignments, of same strands (red) and opposite strands (blue). The vertical stripes indicate repeat annotations in the reference genome: tandem repeats (purple) and transposable elements (pink). (a) Six reads from a BAFME patient that cover the disease-causing *SAMD12* AAAAT repeat locus. (b) Close-ups of three reads with ~5k expansions. (c) Two examples of chimeric human reads (rel3) with expanded CAG repeats at the *ATXN7* disease locus.

### Some difficulties with tandem repeat analysis

The three expanded BAFME reads do not align to the repetitive region as would be expected for a straightforward repeat expansion (Figure 6b). Read 2, from the forward genome strand, does not align to the repetitive region at all, because its expanded region consists mostly of TCCCC repeats whereas the forward strand of the reference genome has TAAAA repeats (Supplemental Figure 2). Reads 1 and 6, from the reverse genome strand, align to the repeat at only one side of the expansion. The expanded regions of these two reads start with TTTTA repeats, which match the reverse strand of the reference, but mostly consist of TTTTC repeats. Since the expanded region of read 2 does not match the reverse complement of read 1 or 6, we infer that systematic sequencing error has occurred on at least one strand. Short-period tandem repeats are prone to a nasty kind of sequencing error: if a systematic error occurs for one repeat unit, the same error will tend to occur for all the other units, producing a different repeat (which may align elsewhere in the genome: the main reason for step 2 in `tandem-genotypes`).

Another kind of difficulty is illustrated by our chimeric human/plasmid reads for *ATXN7* (Figure 6c). Here, the reference sequence adjacent to the annotated repeat is similar to the sequence within the repeat. Depending on the exact sequences and alignment parameters, the expanded region of a read may align outside the repeat annotation (Figure 6c top), or appear as alignment gaps some distance beyond the repeat (Figure 6c bottom). `tandem-genotypes` handles such cases, up to a point, by examining the alignments out to ad hoc distances beyond the annotated repeat.

### Specificity of repeat-expansion predictions

`tandem-genotypes` can handle custom-made repeat annotation files in BED-like format. We made an annotation file with 31 repeat expansion disease loci, including BAFME, and analyzed these 31 repeats with our BAFME data. No large pathological expansions other than BAFME were predicted (Supplemental Figure 3). We also analyzed these 31 repeats with each of the nanopore and PacBio datasets for NA12878: no obvious pathological expansions were predicted and peaks are around zero in most cases (Supplemental Figure 4, 5). These results suggest that our method does not spuriously predict pathological repeat expansions, although there may be some difficulties detecting small disease-causing expansions (e.g. +2 alanine expansion in *PABPN1* causes disease) due to deviations toward copy-number increase in PacBio sequences. We believe this will be solved when sequencing quality improves.

### Prioritization of copy number changes

We also tested if it is possible to prioritize the detected copy number changes. The BAFME repeat expansion was ranked 4^th^ out of 0.7 million tandem repeat regions in rmsk.txt (Figure 7a). When prioritization was done without any control datasets, it was ranked 13^th^, so using controls greatly improved prioritization (Figure 7c).

**Figure 7.**
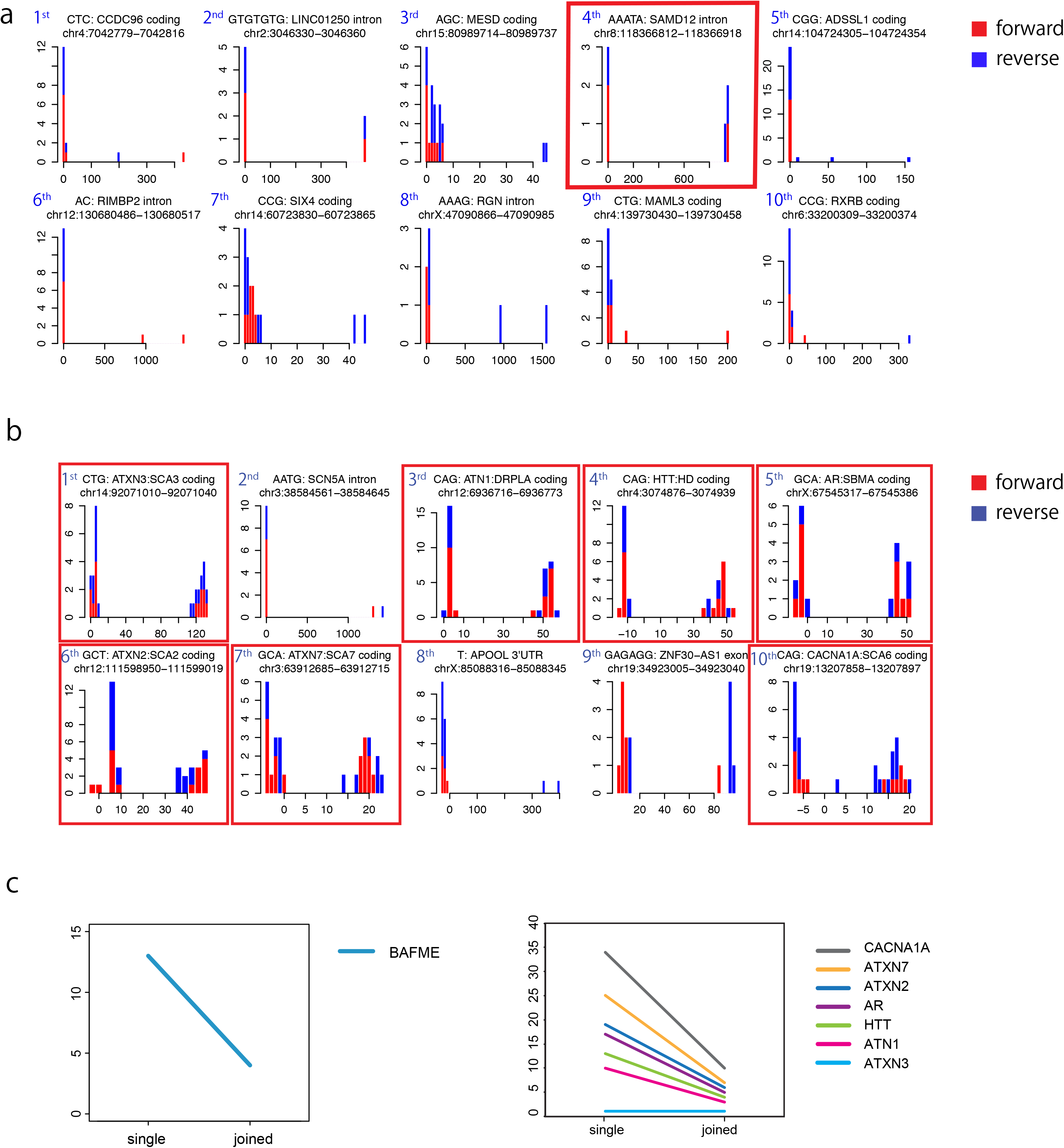
(a) Prioritization of predicted repeat copy number changes in a BAFME patient. The BAFME expansion (AAATA: SAMD12 intron) is ranked 4th out of 0.7 million tandem repeats annotated in rmsk.txt. Forward (red) and reverse strand reads (blue) are shown separately. Histograms are raw output of tandem-genotypes-plot. (b) Prioritization of predicted repeat copy number changes in: whole genome nanopore reads (NA12878 rel3) plus chimeric human/plasmid reads with pathological triplet-repeat expansions (AR, ATN1, ATXN2, ATXN3, ATXN7, CACNA1A and HTT). These pathological expansions are prioritized within the top 10 out of 0.7 million tandem repeats in the genome, when compared to three control datasets using tandem-genotypes-join. (c) Comparison to control datasets with tandem-genotypes-join (joined) effectively de-prioritized other repeats, versus only using a single sample (single). y-axis: prioritization ranking.

Repeat expansions in coding regions can cause disease with less than 100 extra copies, due to the high impact on proteins even from relatively small expansions. So these expansions may be difficult to prioritize. To test this, we combined `tandem-genotypes` output for the whole genome (NA12878 rel3) with outputs for the plasmid-rel3 chimeric reads with coding-region expansions (*ATN1, HTT, ATXN2, ATXN3, CACNA1A, ATXN7* and *AR*). All 7 chimeric expansions were ranked in the top 10 out of 0.7 million repeat regions (Figure 7b). Again, controls greatly improved prioritization (Figure 7c).

### Comparing tandem repeat copy number change distribution of MinION, PromethION and Pacbio RSII sequencing data

We next examined the genome-wide repeat copy number changes in the NA12878 human genome sequenced by both PacBio RSII (SRR3197748) and Oxford Nanopore Technology’s MinION (rel3). There was marked discordance between MinION and Pacbio when the repeat unit size was one or two (Figure 8a, Supplemental Figure 6). This is probably because MinION tends to make small deletion errors and PacBio small insertion errors, which are hard to distinguish from copy number changes of these tiny repeat units. Note that a repeat unit size of one means homopolymers (such as AAAAAAAAAA). The triplet-repeat distribution showed a slight difference between MinION and PacBio (Figure 8a). However, where the repeat unit is longer than 3, MinION and PacBio had similar distributions of copy number changes (Supplemental Figure 6), so both sequencing platforms work on these tandem-repeats. We also verified that the GGGGCC strand bias, which we observed in the plasmids, is also seen in the rel3 dataset. The distribution of GGGGCC copy number change showed a slight difference between forward and reverse strands (Supplemental Figure 1b).

**Figure 8.**
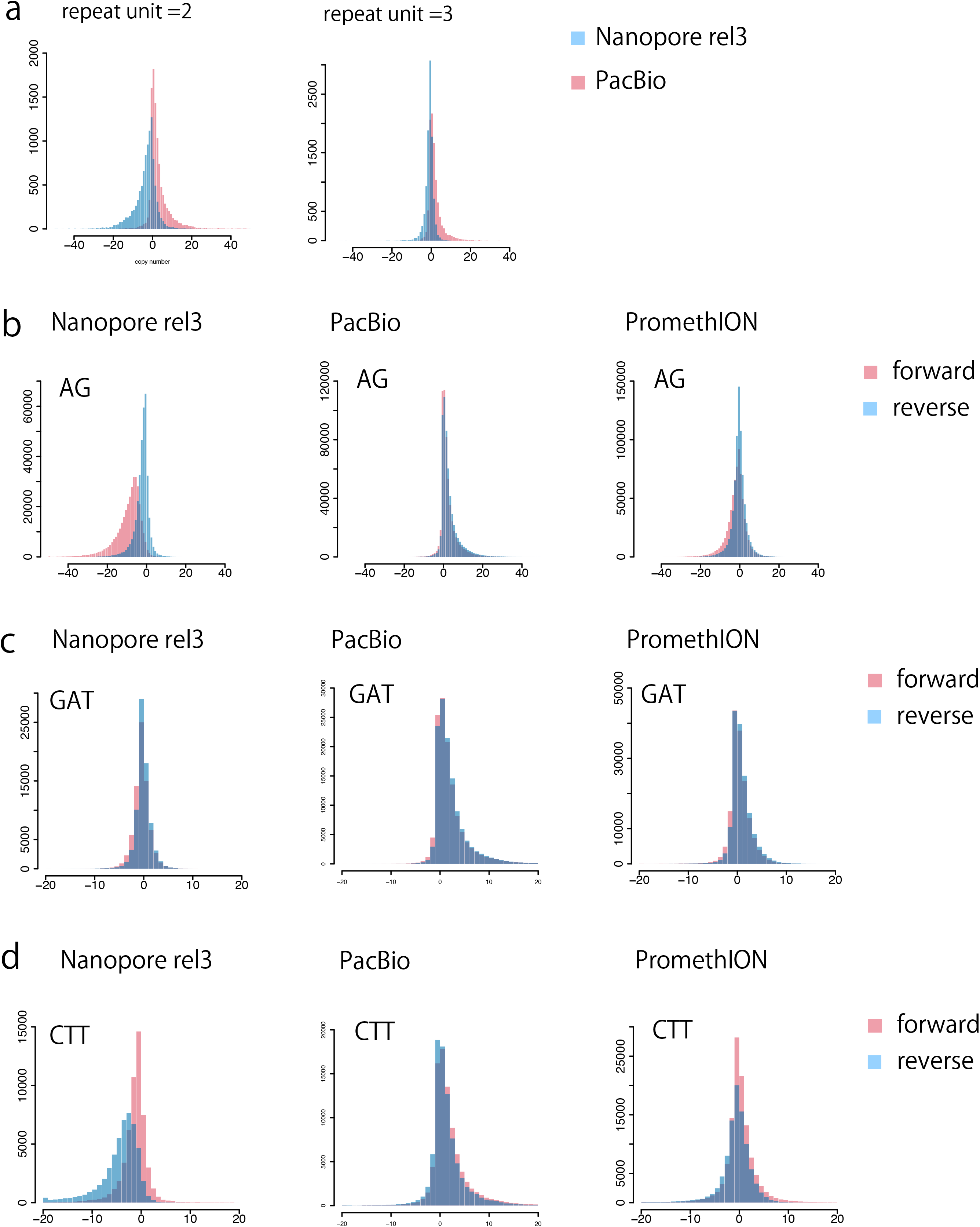
(a) Genome-wide distribution of predicted change in repeat copy number, for nanopore MinION (rel3) and PacBio (SRR3197748) reads from the same human (NA12878). Nanopore tends to have negative and PacBio positive predicted changes, especially for short repeat units. Read number = 10,000 (randomly sampled). Nanopore reads are shown in blue and PacBio in red. (b-d) Genome-wide distribution of predicted change in repeat copy number for nanopore MinION (rel3) and PacBio (SRR3197748) reads from the same human (NA12878), and nanopore PromethION (ERR2585112-5) from a different individual (NA19240). (b) Distributions for AG dinucleotide repeats. (c) Distributions for GAT. (d) Distributions for CTT. CTT shows the most prominent strand bias in nanopore rel3 reads among all types of triplet repeat (all types are in Supplemental Figures 8-13). PromethION shows less strand bias compared to rel3. y-axis: read count, x-axis: change in copy number relative to the reference human genome.

We examined the distribution of copy number change for all kinds of di-nucleotide and triplet repeat. There are four possible types of di-nucleotide repeat (AG, AT, CG and AC) and ten kinds of triplet repeat (TAA, GTC, AAC, GAT, CTT, CTG, GTA, CGG, CCT and CAC). MinION sequences (rel3) showed obvious systematic strand biases for AC, AG, CCT and CTT (Figure 8b,d, Supplemental Figure 7-9). Note that we do not expect any strand bias for CG and AT because they are palindromic.

The MinION data we tested (rel3) was published in 2017. Nanopore base-callers and chemistries have been greatly improved recently. We tested a recent nanopore MinION dataset analyzed by MinKNOW1.11.5, which uses the Albacore 2.0 base-caller. The strand bias for AC, AG, CCT and CTT was greatly improved (Supplemental Figure 10). We also tested a recently published human genome dataset sequenced by ONT’s new high throughput sequencer PromethION. We also found that strand biases for AC, AG, CTT and CCT are greatly improved in PromethION reads (Figure 8b,d, Supplemental Figures 11, 12).

### Computation time and repeat-masking

The slowest computational step was aligning the reads to the genome (`lastal`). For some datasets, we made it much faster by “masking” repeats (both interspersed and tandem) with `windowMasker`. Here, “masking” means that the repeats (indicated by lowercase letters) are excluded from the similarity-search steps of the alignment algorithm, but are included when making the final alignments: the hope is to find the same alignments faster.

In practice, this masking is often harmless, but sometimes harmful. It did not prevent us from detecting the BAFME expansion, or expansions at 10 other disease loci in chimeric human/plasmid reads. On the other hand, it prevented detection of the SCA10 expansions (result not shown). This is because one flank of the SCA10 tandem repeat consists of transposable elements and is almost completely masked. Note this dataset has somewhat short reads (Table 2): the problem would be solved by longer reads that extend beyond the masked region.

When we do not mask, the total run time for our analysis is competitive with those of RepeatHMM and PacmonSTR (Supplemental Table 3). (Note the last-train run time does not increase much for larger datasets, because it uses a fixed-size sample of the data.) When we do mask, the computation is much faster (Supplemental Table 4), and usually the results do not change significantly (Supplemental Figure 13).

## Discussion

We have presented several lines of evidence that we can robustly detect pathological expansions of tandem repeats. We successfully detected them in: constructed plasmids, semi-artificial plasmid/human sequences, and real human sequences from patients with SCA10 and BAFME. We also did *not* detect unexpected (false-positive) large known-pathological expansions in three whole-genome datasets: PacBio reads from a BAFME patient, and PacBio and nanopore reads from NA12878. Importantly, we can also rank copy-number changes by priority, such that pathological expansions are ranked near the top out of ~0.7 million tandem repeats in the genome.

Our method is not specific to tandem expansions, but detects any kind of expansion of a tandem repeat. For example, we detected an expansion due to insertion of an Alu SINE within a tandem repeat (Figure 5a). We also detected an expansion that contained non-repeat sequence, which turned out to be a deletion in the reference genome (Figure 5b). Such non-tandem expansions may impact genomic function and health, so we believe it is useful to detect them too during first-round genome-wide screening.

If a repeat expansion is actually the ancestral state, with the reference genome having a contraction or deletion (e.g. Figure 5b), then it is plausible that the expansion is less likely to be pathological. Thus, our prioritization of copy-number changes likely benefits from comparing the changes to ape genomes. An ancestral reference human genome would be ideal^14^. A similar idea is to de-prioritize expansions commonly present in healthy humans (Figure 7a, b): this will become more powerful as tandem repeat data accumulates.

We have also pointed out some interesting difficulties with analyzing tandem repeat sequences. Some DNA reads do not align to the repetitive region of the reference genome (e.g. Figure 6b), and systematic sequencing errors may turn a tandem repeat into a different tandem repeat. The analysis becomes harder when the reference sequence next to an annotated tandem repeat resembles the sequence in the repeat (e.g. Figure 6c). Some (inexact) tandem repeats do not have unambiguous boundaries, and different annotations (e.g. RepeatMasker versus Tandem Repeats Finder^25^) sometimes disagree on the boundaries. In some cases, there may be no unambiguous distinction between expansion of a tandem repeat and sequence insertion near the repeat.

Systematic sequencing errors can have different effects on the two strands of tandemly-repeated DNA, causing the predicted copy number changes to have a bimodal distribution (e.g. Figure 1i). So it is important to indicate which predictions come from which strands, in order to not misinterpret this as two alleles. We report length and strand biases of several long-read sequencers for every possible type of di- and tri-nucleotide repeat: these biases are prominent for specific repeats (e.g. CTT and CCT in older MinION data), and the worst biases are greatly improved in more recent sequencing systems.

If sequencing accuracy continues to improve, tandem repeat analysis will obviously benefit. The alignment will automatically become faster, due to lower tolerance of gaps and substitutions. Copy number will be predictable more accurately and with lower coverage.

This study demonstrates a practical and robust way to identify changes in tandem repeats that may have biologically impactful consequences. Although there are still limitations due to the developing sequencing technologies and cost to immediately apply this approach in clinical sequencing, we clearly show that there is hope that long read sequencing is useful to identify overlooked changes in the genome, and may give an answer to the large numbers of patients with genetic diseases whose causes and mechanisms have remained unsolved for many years.

## Acknowledgements

MCF thanks Masahiro Onoguchi for pointing out that strand bias must be discriminated from heterozygosity. SM thanks to Qian Liu for suggesting parameters for SCA10 data analyses using RepeatHMM. YK thanks Jun-ichi Satoh and Mika Takitani for plasmid construction. Sanger sequencing was performed by the Support Center for Medical Research and Education, Tokai University.

## Author contributions

SM and MCF contributed to the conception of the work, acquisition, analysis and interpretation of the data T.M, S.M, T.T, H.A, Y.O, Y.K, H.M and N.M contributed to obtaining and making the materials and acquisition and analysis of data.

## Reference

1. Tang H, Kirkness EF, Lippert C, et al. Profiling of Short-Tandem-Repeat Disease Alleles in 12,632 Human Whole Genomes. Am J Hum Genet 2017;101:700–15.

2. La Spada AR, Roling DB, Harding AE, et al. Meiotic stability and genotype-phenotype correlation of the trinucleotide repeat in X-linked spinal and bulbar muscular atrophy. Nat Genet 1992;2:301–4.

3. A novel gene containing a trinucleotide repeat that is expanded and unstable on Huntington’s disease chromosomes. The Huntington’s Disease Collaborative Research Group. Cell 1993;72:971–83.

4. Brook JD, McCurrach ME, Harley HG, et al. Molecular basis of myotonic dystrophy: expansion of a trinucleotide (CTG) repeat at the 3’ end of a transcript encoding a protein kinase family member. Cell 1992;68:799–808.

5. Kremer EJ, Pritchard M, Lynch M, et al. Mapping of DNA instability at the fragile X to a trinucleotide repeat sequence p(CCG)n. Science 1991;252:1711–4.

6. Lemmers RJ, van der Vliet PJ, Klooster R, et al. A unifying genetic model for facioscapulohumeral muscular dystrophy. Science 2010;329:1650–3.

7. Brais B, Bouchard JP, Xie YG, et al. Short GCG expansions in the PABP2 gene cause oculopharyngeal muscular dystrophy. Nat Genet 1998;18:164–7.

8. Musova Z, Mazanec R, Krepelova A, et al. Highly unstable sequence interruptions of the CTG repeat in the myotonic dystrophy gene. Am J Med Genet A 2009; 149A: 1365–74.

9. McFarland KN, Liu J, Landrian I, et al. SMRT Sequencing of Long Tandem Nucleotide Repeats in SCA10 Reveals Unique Insight of Repeat Expansion Structure. PLoS One 2015;10:e0135906.

10. Ishiura H, Doi K, Mitsui J, et al. Expansions of intronic TTTCA and TTTTA repeats in benign adult familial myoclonic epilepsy. Nat Genet 2018;50:581–90.

11. Nishikawa A, Mitsuhashi S, Miyata N, Nishino I. Targeted massively parallel sequencing and histological assessment of skeletal muscles for the molecular diagnosis of inherited muscle disorders. J Med Genet 2017;54:104–10.

12. Ummat A, Bashir A. Resolving complex tandem repeats with long reads. Bioinformatics 2014;30:3491–8.

13. Liu Q, Zhang P, Wang D, Gu W, Wang K. Interrogating the “unsequenceable” genomic trinucleotide repeat disorders by long-read sequencing. Genome Med 2017;9:65.

14. Frith MC, Khan S. A survey of localized sequence rearrangements in human DNA. Nucleic Acids Res 2018;46:1661–73.

15. Hamada M, Ono Y, Asai K, Frith MC. Training alignment parameters for arbitrary sequencers with LAST-TRAIN. Bioinformatics 2017;33:926–8.

16. Frith MC. A new repeat-masking method enables specific detection of homologous sequences. Nucleic Acids Res 2011;39:e23.

17. Frith MC. Gentle masking of low-complexity sequences improves homology search. PLoS One 2011;6:e28819.

18. Jain M, Koren S, Miga KH, et al. Nanopore sequencing and assembly of a human genome with ultra-long reads. Nat Biotechnol 2018;36:338–45.

19. Oma Y, Kino Y, Sasagawa N, Ishiura S. Intracellular localization of homopolymeric amino acid-containing proteins expressed in mammalian cells. J Biol Chem 2004;279:21217–22.

20. Kino Y, Washizu C, Kurosawa M, et al. Nuclear localization of MBNL1: splicing-mediated autoregulation and repression of repeat-derived aberrant proteins. Hum Mol Genet 2015;24:740–56.

21. Oma Y, Kino Y, Toriumi K, Sasagawa N, Ishiura S. Interactions between homopolymeric amino acids (HPAAs). Protein Sci 2007;16:2195–204.

22. O’Leary NA, Wright MW, Brister JR, et al. Reference sequence (RefSeq) database at NCBI: current status, taxonomic expansion, and functional annotation. Nucleic Acids Res 2016;44:D733–45.

23. Morgulis A, Gertz EM, Schaffer AA, Agarwala R. WindowMasker: window-based masker for sequenced genomes. Bioinformatics 2006;22:134–41.

24. Nakamura K, Oshima T, Morimoto T, et al. Sequence-specific error profile of Illumina sequencers. Nucleic Acids Res 2011;39:e90.

25. Benson G. Tandem repeats finder: a program to analyze DNA sequences. Nucleic Acids Res 1999;27:573–80.

26. Edgar RC. MUSCLE: multiple sequence alignment with high accuracy and high throughput. Nucleic Acids Res 2004;32:1792–7.

